# Rapid adaptive optics enabling near noninvasive high-resolution brain imaging in awake behaving mice

**DOI:** 10.1101/2025.05.26.656230

**Authors:** Zhentao She, Yiming Fu, Yingzhu He, Gewei Yan, Wanjie Wu, Zhongya Qin, Jianan Qu

**Affiliations:** Department of Electronic and Computer Engineering and State Key Laboratory of Molecular Neuroscience, The Hong Kong University of Science and Technology, Hong Kong, P. R. China

**Author notes:** These authors contributed equally: Zhentao She, Yiming Fu.

## Abstract

High-resolution imaging under physiological conditions is essential for studying biological mechanisms and disease processes. However, achieving this goal remains challenging due to optical aberrations and scattering from heterogeneous tissue structures, compounded by motion artifacts from awake animals. In this study, we developed a rapid and accurate adaptive optics system called multiplexing digital focus sensing and shaping (MD-FSS) for deep-tissue multiphoton microscopy. Under two-photon excitation, MD-FSS precisely measures the aberrated point spread function in approximately 0.1 s per measurement, effectively compensating for both aberrations and scattering to achieve subcellular resolution in deep tissue. Using MD-FSS integrated with two-photon microscopy, we achieved high-resolution brain imaging through thinned or optically cleared skull windows, two near noninvasive methods to access mouse brain, reaching depths up to 600 μm below the pia in awake behaving mice. Our findings revealed significant differences in microglial functional states and microvascular circulation dynamics between awake and anesthetized conditions, highlighting the importance of studying brain function in awake mice through noninvasive methods. We captured functional imaging of fine neuronal structures at subcellular level in both somatosensory and visual cortices. Additionally, we demonstrated high-resolution imaging of microvascular structures and neurovascular coupling across multiple cortical regions and depths in the awake brain. Our work shows that MD-FSS robustly corrects tissue-induced aberrations and scattering through rapid PSF measurements, enabling near-noninvasive, high-resolution imaging in awake, behaving mice.

## Introduction

Advances in microscopy technologies for biological imaging have significantly enhanced our understanding of physiological processes. Multiphoton microscopy (MPM) has emerged as a powerful tool, leveraging nonlinear excitation to enable *in vivo* imaging in living animal models for studying fundamental biology and disease mechanisms^1–3^. However, tissue heterogeneity induces aberrations and scattering that distort the excitation light wavefront, limiting both resolution and depth in noninvasive deep-tissue imaging. In recent years, adaptive optics (AO), originally developed in astronomy to correct atmospheric aberrations, has been successfully adapted to microscopy to correct tissue-induced aberrations and improve both imaging resolution and depth^4–19^.

AO methods can be classified into two approaches based on aberration measurement: indirect and direct wavefront sensing. Indirect methods use algorithmic optimization of correction wavefronts based on fluorescence signal metrics^7–12^, while direct methods employ dedicated wavefront sensors for aberration measurement and correction^13–16^. However, the limited speed of traditional AO methods restricts their application to anesthetized animals, despite evidence that anesthesia alters normal physiological states and cellular behavior^20–25^. While imaging awake animals would provide more accurate biological insights, their increased movement compared to anesthetized subjects generates significant motion artifacts^26–31^. These motion artifacts compromise AO measurement accuracy by disrupting the stable guidestar required for wavefront correction. Therefore, achieving precise wavefront correction for near noninvasive MPM in awake, behaving animals requires an AO method that combines both speed and accuracy.

We recently reported an AO technology called ALPHA-FSS for three-photon microscopy (3PM) that directly measures the aberrated electric-field point spread function (*E-PSF*) through raster scanning between a strong beam and a coherent weak beam, with aberration correction achieved through phase conjugation of the measured PSF^17^. This technology shares the same physical principle as F-SHARP^18,19^. ALPHA-FSS demonstrated successful probing and correction of both aberrations and scattering *in vivo* through highly turbid tissues, including intact mouse skull. However, despite its advantages of high accuracy, high correction order, and relatively fast measurement speed, the several-second duration required for PSF measurement remains a limiting factor for applications in awake behaving animals due to motion artifacts. The PSF measurement duration is largely limited by the low repetition rate of the three-photon laser and the low efficiency of three-photon excitation (3PE). Due to its higher-order nonlinear excitation, 3PE generates significantly fewer fluorescence photons compared to two-photon excitation (2PE) at the same excitation intensity and pulse energy^32,33^. This necessitates a longer dwell time at each PSF sampling location during the raster scanning process on a three-photon microscope. Therefore, 2PE is more advantageous for rapid PSF measurement though it offers less penetration capability in tissue^18,34^.

In this study, we developed an AO method called multiplexing digital focus sensing and shaping (MD-FSS), combined with a two-photon microscopy (2PM), to achieve rapid AO measurement and correction for high-resolution brain imaging in awake mice. Technically, MD-FSS consists of two major innovations: multi-beam interference for fast focal field sensing and digital phase demodulation for parallel measurement of PSF’s complex amplitude. In brief, MD-FSS operates on a similar principle of aberration measurement as ALPHA-FSS^17^, but remarkably accelerates the PSF measurement speed by simultaneously interfering multiple modulated weak beams, each with small spatial and frequency offsets, with a strong beam. The system detects the phase and amplitude of the modulated interference signals at their corresponding modulation frequencies using multi-channel digital Fast Fourier Transform (FFT) detection. The detected down-sampled *E-PSF* from each weak scanning beam is then combined to reconstruct a full-sampled PSF without compromising accuracy, enabling rapid and precise wavefront correction through phase conjugation.

When applied to two-photon brain imaging in awake behaving mice, MD-FSS enables accurate measurement of the aberrated PSF that is robust against motion artifacts. Through thinned and optically cleared skull windows, methods for near noninvasive optical access to the mouse brain, it achieves subcellular resolution at depth. We demonstrated MD-FSS-2PM performance through *in vivo* subcellular imaging of microglial and microvascular dynamics in the awake mouse brain, revealing distinct morphological and functional differences between awake and anesthetized states. Furthermore, we achieved high-resolution functional imaging of neurons in both somatosensory and visual cortices at depth, with calcium transients in apical and basal dendrites clearly visible after AO correction with MD-FSS. Additionally, we captured neurovascular coupling through simultaneous imaging of neuronal activity and blood vessels in multiple cortical regions of the awake behaving mouse following AO correction.

## Results

### Rapid Complex-valued *E-PSF* Sensing via MD-FSS

Briefly, MD-FSS reduces the measurement time for PSF by using beam multiplexing and multi-channel digital FFT demodulation. To measure the aberrated PSF in deep tissue, multiple weak beams with different modulation frequencies and spatial shifts are generated by an acousto-optic deflector (AOD) to interfere with a strong beam in the focal plane (Fig. 1a, Fig. S1). The multiple weak beams are then raster-scanned relative to the strong beam simultaneously, and the 2P fluorescence intensity *I* at the scanning position *x* is given by:

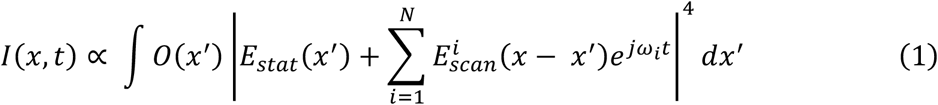

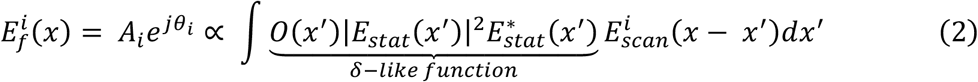

**Fig. 1.**
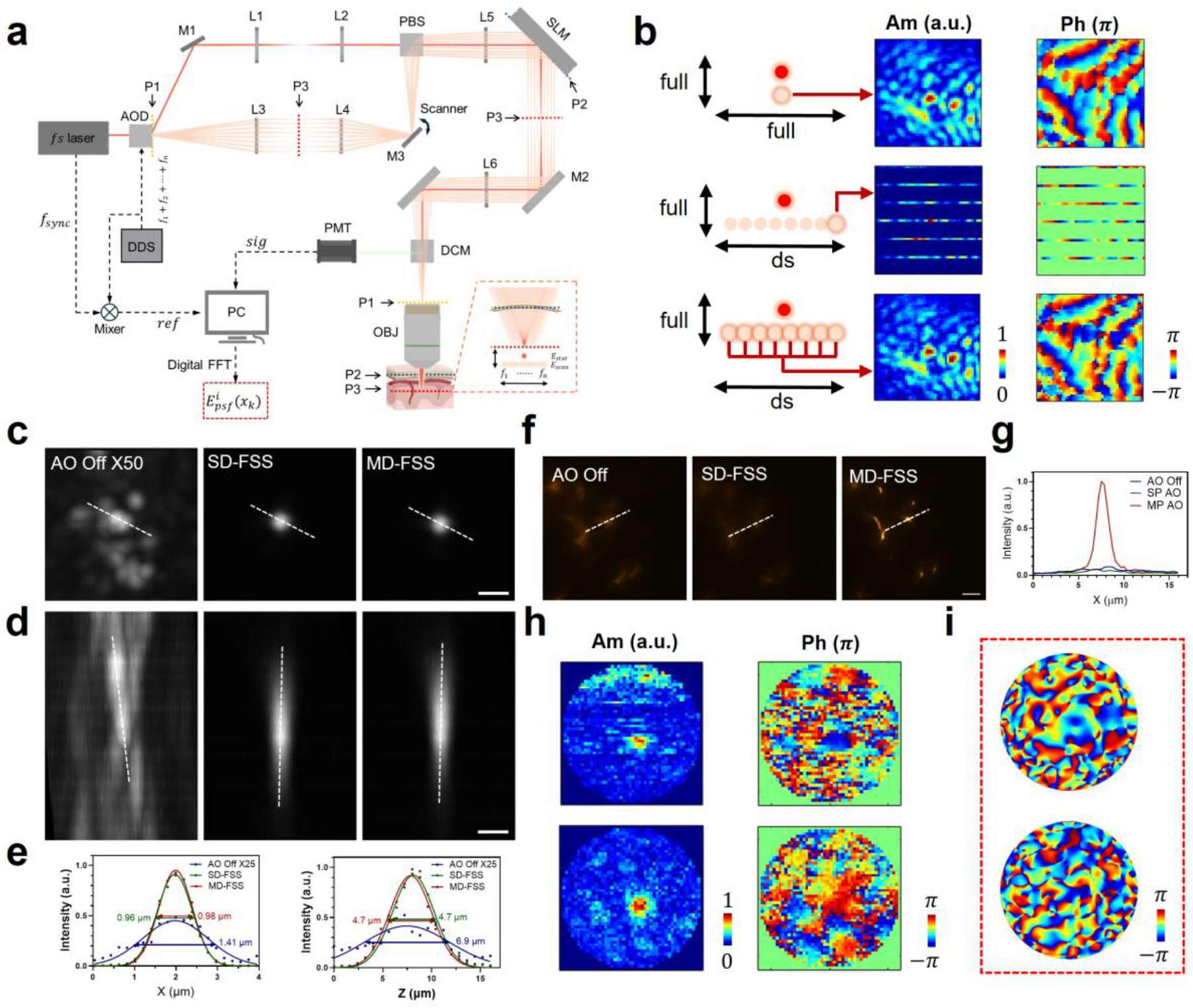
MD-FSS enables rapid aberration measurement for AO correction through highly turbid medium. (a) Schematics of MD-FSS system. Inset illustrates the measurement of aberrated PSF electric field by MD-FSS: Multiple phase-modulated weak focal points are spatially shifted by equal distances to simultaneously probe the electric field. *E*_*stat*_, focal field of strong stationary beam positioned on the guide star; *E*_*scan*_, focal field of the multiple weak scanning beams; 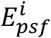, measured complex-valued PSF of *i*_*th*_ multiplexed beam at down-sampled scanning scheme; *f*_*n*_, modulation frequency of each weak scanning beam. (b) Comparison of the measured PSFs using SD-FSS and MD-FSS. Top: Diagram illustrating the focal field and scanning scheme of SD-FSS (left), accompanied by the amplitude (mid) and phase (right) of the full-sampled PSF measured by SD-FSS; Middle: Diagram illustrating the scanning scheme of one down-sampled scanning beam in MD-FSS (left), along with the amplitude (mid) and phase (right) of one down-sampled PSF; Bottom: Diagram illustrating the merging of multiple down-sampled PSFs from all scanning beams (left), along with the amplitude (mid) and phase (right) of the merged full-sampled PSF measured by MD-FSS; ds: down-sampled scanning, full: full-sampled scanning. (c, d) 2PF image of 1 *μm* fluorescence beads at a depth of 400 *μm* through a 50 *μm* thinned skull, shown without AO correction (left), with SD-FSS AO correction (middle) and with MD-FSS correction (right); Signal intensity with AO correction was enhanced by 50-fold. (e) Line profile and full width at half maximum (FWHM) measured along the dashed white line in (c) (left) and (d) (right). (f) Comparison of AO correction result between SD-FSS and MD-FSS on awake behaving Thy1-YFP mouse through a thinned-skull window; Images of neuron dendrites without AO correction (left), with SD-FSS AO correction (middle), and with MD-FSS AO correction (right). (g) Line profile along the dashed white line in (f). (h) Measured amplitude and phase of aberrated PSF using SD-FSS (top) and MD-FSS (bottom) in (f) on awake behaving mouse; (i) AO correction phase pattern calculated from PSF in (h), using SD-FSS (top) and MD-FSS (bottom). The results (f)-(i) demonstrate that MD-FSS’s fast measurement speed enables robust performance against motion artifacts. Scale bars: 1 *μm* in (c, d); 20 *μm* in (f).

Here, *O*(*x*^′^) represents the real-valued object function, *ω*_*i*_ denotes the modulation frequency of each scanning beam, *N* is the total number of weak scanning beams, *E*_*stat*_(*x*^′^) is the complex-valued focal field of the strong stationary beam, and 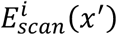 is the complex-valued focal field of the *i*_*th*_ weak scanning beam. By setting the intensity of each scanning beam to be much less than that of the stationary beam 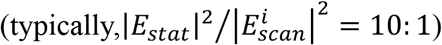, the oscillating components of the 2P fluorescent intensity *I* with different modulation frequencies {*ω*_*i*_}, can be demodulated using digital FFT detection (See Methods, Fig. S2). The digital FFT method requires only recording of the modulated fluorescence signal and reference, allowing for the simultaneous output of amplitude and phase components *A*_*i*_ and *θ*_*i*_ for focal field 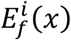 of each weak scanning beam through digital calculation, significantly simplifying the process of multi-frequency demodulation.

The complete aberrated PSF can be reconstructed from the down-sampled PSF of each scanning beam (Fig. 1b), and the value of the restored complete PSF 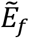 at position *x*_*k*_ can be calculated using the following formula:

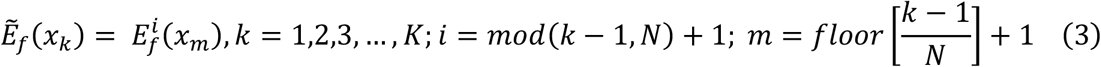

Here, *K* represents the pixel number of the full-sampled PSF, *mod* denotes the modulo operator and *floor* represents rounding to nearest integer less than the element. With the reconstructed full-sampled PSF, the correction wavefront is generated through phase conjugation by performing the 2D FFT^17^.

The principle of MD-FSS involves generating multiple weak scanning beams with different phase modulation frequencies and controllable spatial shifts in the focal plane. In this work, we developed a straightforward method to generate the desired multiple scanning beams using an AOD, which diffracts the input beam with applied radio frequency (RF) voltage that determines the frequency and angle of the diffracted beam (see Methods and Fig. S3). By applying an RF signal with multiple frequencies to the AOD, the light is diffracted into several beams, each with unique frequencies and spatial positions, as determined by the multi-frequency RF signal. In matching the down-sampled scanning scheme, we chose the RF modulation frequencies, ensuring the spatial shift between adjacent scanning beams in the focal plane equals the line distance of the full-sampled scanning scheme. Since AOD is a highly dispersive component, the focus of the diffraction beam is elongated by the angular dispersion, compromising the quality of multi-beam interference. To compensate for the angular dispersion, a single prism-AOD configuration^35^ was previously used to match the dispersion by adjusting the incidence angle, while another method employing a second AOD^36^ provided a controllable dispersion value with minimal beam expansion and power loss, though it requires more complex control. For the dispersion compensation in this application, we developed a simpler dual prism configuration method that eliminates power loss and extra complexity, making it both effective and easy to adjust (see Methods and Fig. S4). In this work, we multiplexed eight weak scanning beams to measure the aberrated PSF, accelerating the process by up to eight times and reducing the measuring time to approximately 0.1 s for one PSF.

With the measured down-sampled PSFs by each of the eight scanning beams, it is essential to calibrate the relative amplitude and phase offsets between adjacent beams for accurate reconstruction of a full-sampled PSF. We developed a calibration method for accurately measuring the relative amplitude and phase (see Methods), where the PSFs of each weak beam are measured simultaneously using fluorescence beads as the guidestar without down-sampling. The amplitude ratio lookup table (LUT) is calculated by dividing the mean amplitude of each PSF by that of the first beam’s PSF, while the phase offset LUT is determined by sequentially adjusting the phase map of each PSF and comparing it to the first beam’s phase map, recording the value that yields the best correlation as the correct phase offset for each scanning beam.

To assess the MD-FSS method for measuring the complex-valued *E-PSF* and correcting aberrations in a turbid medium, we performed an *in-vitro* experiment imaging stable fluorescent beads through a thinned skull extracted from an adult mouse. Using the fluorescent beads as a guidestar, we simultaneously measured the down-sampled PSF of each scanning beam and merged them, with calibrated relative amplitude and phase offsets, into a full-sampled PSF. For comparison, we used the full-sampled PSF measured with single beam digital focus sensing and shaping (SD-FSS) as the reference PSF. SD-FSS employs a single scanning beam to interfere with a stationary beam, equivalent to our previously reported ALPHA-FSS^17^, providing optimal SNR for a stable guide star (Fig. S5). We found that the merged PSF from MD-FSS maintained the same SNR as the reference PSF (Fig. 1b, S6a, S7) at the same intensity of the strong beam. Next, we verified MD-FSS performance in correcting skull-induced aberrations compared to SD-FSS (Fig. 1c-e, S6). Before AO correction, the PSF was severely distorted into numerous small side lobes due to the highly turbid skull. After AO correction using MD-FSS, the signal intensity increased by 50-fold, restoring the PSF to near diffraction-limited quality without side lobes. With remote focusing and the conjugation between the spatial light modulator (SLM) and the skull layer^17^, the correction area achieved a field of view (FOV) of approximately 150 *μm* in the lateral direction and 300 *μm* in the axial direction (Fig. S6e, f). Quantitatively, MD-FSS matched SD-FSS in improvements to resolution and signal intensity, while requiring only one-eighth of the measurement time, significantly accelerating the AO measurement process.

With the *in-vitro* experiment showcasing MD-FSS’s performance on a stable fixed sample, we next evaluated its capabilities in measuring the aberrated PSF through thinned skull in awake, behaving mice, where motion artifacts are significant. The thinned skull thickness was maintained above 30 μm to ensure mechanical stability of the cranial window while maintaining a noninvasive approach to brain imaging. We prepared the brain imaging window by securely gluing a metallic headplate to the skull using dental cement and fixing it to a holder with metal screws. Despite careful preparation to prevent looseness, the mouse’s movements caused frequent rapid shifts within the imaging FOV, consistent with previous findings^26–30^ (Supplementary Video S1, 2). MD-FSS’s fast AO measurement speed enabled accurate measurement of the complex-valued aberrated PSF in awake behaving mice, with the derived correction wavefront significantly enhancing both fluorescence intensity and imaging resolution. In contrast, SD-FSS’s longer measurement time led to motion-deteriorated PSF, resulting in a correction wavefront that failed to enhance signal intensity and even introduced additional aberrations degrading imaging quality (Fig. 1f-i, S8, Supplementary Video S1, 2). The results revealed that SD-FSS’s PSF measurement errors in awake, behaving mice primarily arose from off-target motion of the guide star relative to the strong stationary focus, causing fluorescence loss during raster scanning interference signal detection. Under anesthesia, the cessation of motion allowed both MD-FSS and SD-FSS to accurately measure the aberrated PSF, remarkably improving signal intensity and imaging resolution (Fig. S8), proving that MD-FSS is essential for *in vivo* brain imaging of awake mice. In the following section, we present MD-FSS applications for high-resolution imaging of cell morphology and function in the awake behaving mouse brain, achieved in a near noninvasive manner.

It should be pointed out that although the principle of MD-FSS is theoretically applicable to three-photon microscopy, its practical use for rapid AO measurement is limited by the low repetition rate, low nonlinear absorption coefficient, and significantly higher water absorption of the 3P laser^32,33^. We experimentally verified that due to the weak fluorescence intensity of 3PE, it inherently requires a longer dwell time to achieve the same quality of measured PSF as 2PE for AO correction^17,34^. In this study, we focus on implementing MD-FSS-2PM and show its effectiveness for rapid measurement of aberrated *E-PSF*, as well as for correcting both aberrations and scattering *in-vitro* and *in-vivo* in deep tissue imaging.

### MD-FSS Enables Near Noninvasive Microglial Imaging in Awake Behaving Mice

Having demonstrated the unique capabilities of MD-FSS for rapid and accurate aberration measurement through theoretical analysis and experiments, we investigated its performance *in vivo* for structural and functional imaging of microglia in awake, behaving Cx3Cr1-GFP mice through thinned skull (Fig. 2, S9-12). Microglia, the major immune cells in the central nervous system (CNS), are highly dynamic and sensitive to their surrounding physiological environment^37,38^. While mice were freely moving on a treadmill, we measured the aberrated PSF in deep tissue using MD-FSS and applied corresponding corrections (Fig. 2b-e). The results show that before AO correction, aberrations severely blurred the fine structures of microglia, revealing only indistinct soma, but after correction, microglial processes became clearly visible with significantly enhanced signal intensity (Fig. 2b-d, S9-11). To investigate microglial morphology and physiological dynamics in their natural state, we performed time-lapse imaging with AO correction in awake, behaving mice, then anesthetized the mice and repeated the imaging with AO correction (Fig. 2g-k, S9). The high-resolution images enabled detailed analysis of microglial morphological characteristics in both states. We found that under anesthesia, microglia became more ramified, with processes expanding upon isoflurane administration and reaching larger areas compared to the awake state (Fig. 2i-k, S9e, f, Supplementary Video S3). These distinct differences in microglial morphology and behavior between awake and anesthetized states^20,21^ demonstrate that anesthesia, commonly used in *in vivo* imaging, can substantially alter cellular properties of microglia.

**Fig. 2.**
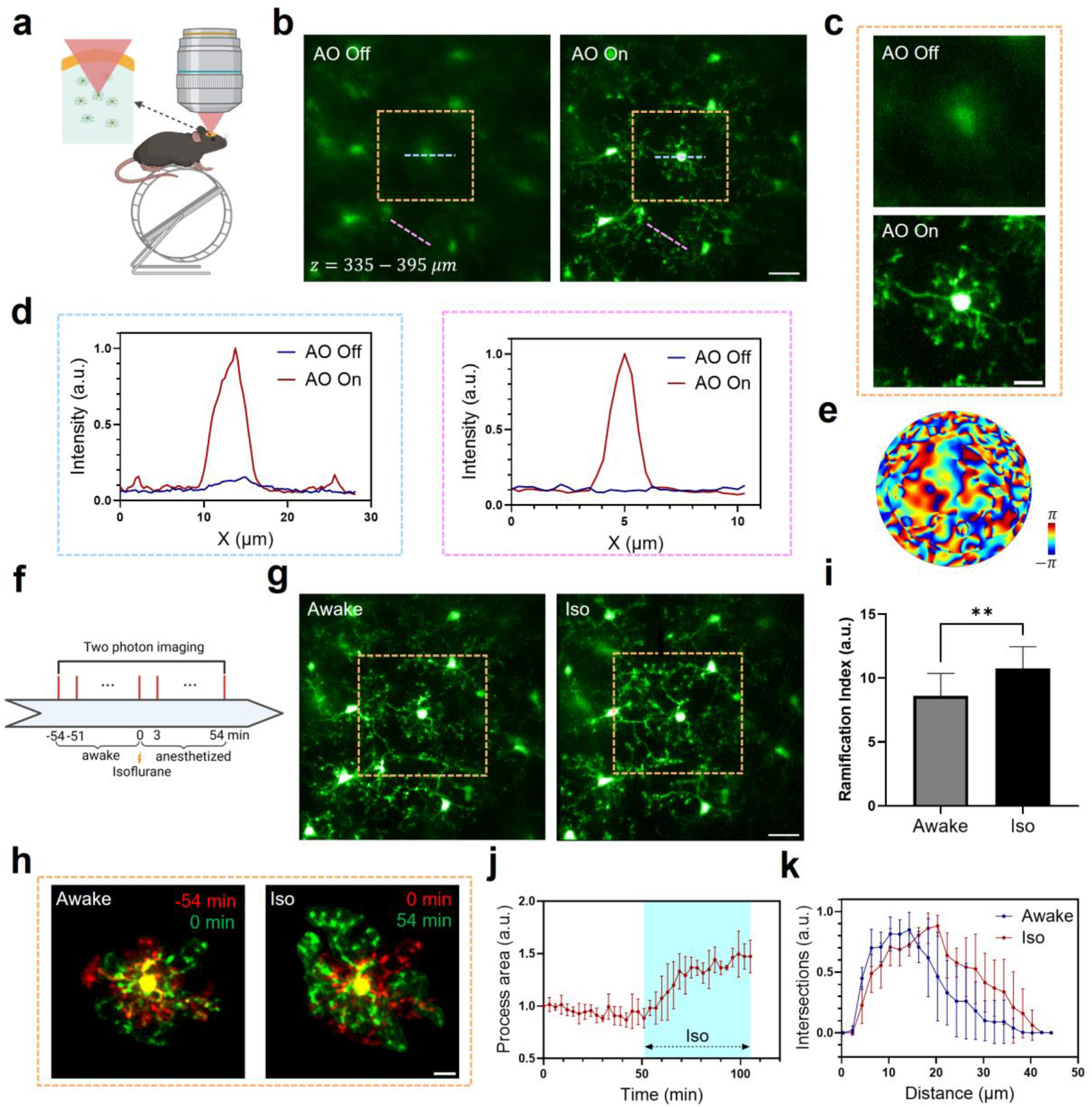
*In-vivo* high-resolution imaging of microglia morphology and dynamics in brain of awake behaving mouse. (a) Schematics of MD-FSS-2PM in brain cortex of awake behaving Cx3Cr1-GFP transgenic mouse. (b) MIP images of GFP-labelled microglia within a volume at a depth of 335 − 395 *μm* below pia, shown without (left) and with (right) MD-FSS AO correction. (c) Zoom-in views of regions in (b) marked by orange dashed box. (d) Intensity profiles along the blue dashed lines (left) and pink dashed lines (right) in (b) without and with AO correction. (e) Correction phase pattern calculated from the PSF measured by MD-FSS. (f) Timeline of MD-FSS-2PM imaging from awake to anesthetized states. (g) MIP images of GFP-labelled microglia in the same volume as (b) from an awake behaving mouse (left) and after 54 min of isoflurane anesthesia (right). (h) Extracted morphological dynamics of microglia in the awake state (left) and anesthetized state (right) within the region indicated by the red dashed rectangle in (f); Red: reference microglial structure 54 minutes before (left) and at 0 min before (right) isoflurane administration; green: microglia morphology at 0 min before (left) and 54 min after (right) isoflurane administration. (i) Statistical analysis of the microglia ramification index in the awake and anesthetized states (Paired *t*-test, *α* = 0.05, N = 5 microglia from 3 mice, **p<0.01, p = 0.0084). (j) Statistical analysis of process area changes before and after isoflurane administration (N = 3 microglia from 3 mice); Blue region: duration of isoflurane administration. (k) Sholl analysis of microglial morphology in awake behaving mouse and anesthetized mouse (N = 4 microglia from 3 mice). Awake: awake state; Iso: anesthetized by isoflurane. Data in (i-k) were represented as mean ± SD. Scale bars: 20 *μm* in (b, g); 10 *μm* in (c, h).

To further investigate microglial dynamics in the awake state, we performed time-lapse imaging following lipopolysaccharide (LPS) injection, which induces global inflammation in the mouse brain^39–42^. The enhanced resolution from AO correction allowed clear distinction of microglial processes and soma, enabling tracking of their dynamic properties after LPS injection. As shown in Fig. S11 and Supplementary Video S4, microglia exhibited activated morphology with significantly reduced process length following LPS injection, consistent with previous observations^40,41^. Through dual-color imaging of microglia and blood vessels, we observed microglial migration toward blood vessels in the awake state after LPS injection (Fig. S12a-f), aligning with previous studies on LPS effects on the blood-brain barrier (BBB)^40^. The high resolution in blood vessel imaging after AO correction enabled precise vessel diameter measurements (Fig. S12g-i), revealing post-LPS increases consistent with previous findings^42^.

Lastly, we performed MD-FSS-2PM imaging through an optical clearing window. Optical clearing is another near noninvasive method for transcranial imaging that chemically increases mouse skull transparency to provide optical access to the brain^43^. Despite increased skull transparency, aberrations from the thick optical clearing skull still severely distort the excitation wavefront, leading to blurred imaging of fine structures in deep tissue. To verify MD-FSS performance with a different optical window, we applied MD-FSS-2PM imaging through an optical clearing window to correct aberrations and restore subcellular resolution in awake, behaving Cx3Cr1-GFP mice. As shown in Fig. S13, microglial processes were clearly visualized after AO correction at different depths, demonstrating that MD-FSS works effectively with both thinned-skull and optical clearing windows. For subsequent studies, we focused on using the thinned-skull window.

### High-Resolution Calcium Imaging of Neuronal Activity in Awake Behaving Mouse

Neuronal activity substantially depends on physiological state and is disrupted by anesthesia^23– 25^, highlighting the need to investigate neurons in awake animals. Calcium imaging, which reflects neuronal activity transients, is a common method for studying neuronal function^44^. We tested MD-FSS-2PM for high-resolution calcium imaging of GCaMP6s-expressing neurons in layers 2/3 and 4 of the somatosensory and visual cortices through thinned skull in awake behaving mice (Fig. 3). In the somatosensory cortex during whisker stimulation, images before AO correction showed blurred soma contours and indistinguishable fine apical dendrites (Fig. 3b). The extracted calcium transients displayed poor signal intensity and low SBR (Fig. 3e). After MD-FSS AO correction (Fig. 3d), neuronal soma borders and lateral dendrites became clearly resolved, with greatly restored apical dendrite details (Fig. 3b, Supplementary Video S5, 6). Calcium transients during whisker stimulation showed significantly improved signal intensity and strong synchronization with stimulation after AO correction (Fig. 3e). Line profiles across identical apical dendrites before and after AO correction revealed substantial resolution enhancement (Fig. 3c). While calcium transients from dendrites could be extracted from AO-corrected images as somas, measurement of these important signals in uncorrected images was impossible due to poor resolution.

**Fig. 3.**
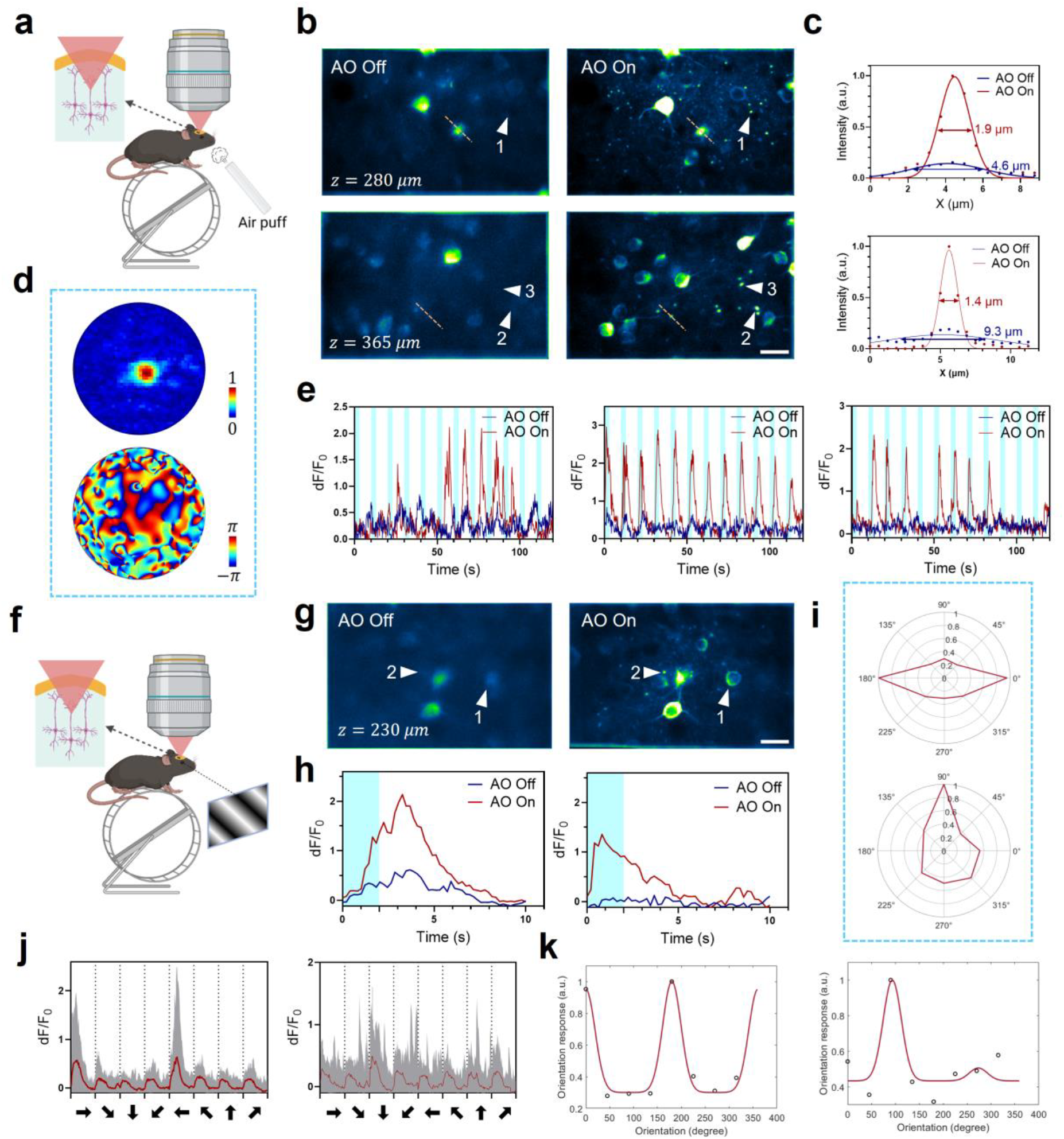
Calcium imaging of neuron somas and dendrites in somatosensory and visual cortices of awake behaving mice. (a) Schematic of MD-FSS-2PM imaging in somatosensory cortex during whisker stimulation in awake behaving CaMKII-GCaMP6s transgenic mice. (b) Standard deviation (STD) projection images of GCaMP6s-labeled neurons recorded over 120 s at 280 *μm* (top) and 365 *μm* (bottom) below pia, without (left) and with (right) AO correction. (c) Intensity profiles along orange dashed lines in (b) at 280 *μm* (top) and 365 *μm* (bottom). (d) PSF amplitude (top) and AO correction phase pattern (bottom) in (b). (e) Calcium transients from neuronal dendrites marked by red arrowheads 1 (left), 2 (middle) and 3 (right) in (b). Blue bars: whisker stimulation periods. (f) Schematic of MD-FSS-2PM imaging in visual cortex during visual stimulation in awake behaving CaMKII-GCaMP6s transgenic mice. (g) STD projection images of GCaMP6s-labeled neurons recorded over 160 s at 230 *μm* below pia, without (left) and with (right) AO correction. (h) Calcium transients at preferred grating orientation from dendrites and soma marked by arrowheads 1 (left) and 2 (right), without and with AO correction. Blue bars: visual stimulation periods. (i) Orientation polar maps of dendrites and soma marked by arrowheads 1 (top) and 2 (bottom) after AO correction. (j) Calcium transients from dendrites and soma marked by arrowheads 1 (left) and 2 (right) during visual stimulation at different orientations with AO correction. Gray curves: raw calcium transients from multiple trials (N = 10) of random orientations; red curves: averaged calcium transients; gray dashed lines: visual stimulation timestamps; black arrowheads: stimulation orientations. (k) Orientation tuning curves with Gaussian fits from regions marked by arrowheads 1 (left) and 2 (right) in (g) after AO correction. Scale bars: 20 *μm* in (b, g).

Using the inertia-free remote focusing module for depth adjustment and the extensive axial correction range of conjugate AO^17,45^, we achieved quasi-simultaneous multi-plane neuronal imaging by rapidly switching the electrically tunable lens (ETL) defocus with synchronized deformable mirror correction patterns (Fig. S14, 15, Supplementary Video S7, 8). Multi-plane imaging enabled investigation of correlated neuronal activity along the axial direction^46–49^. As shown in Fig. S14 and S15, without AO correction, fine structures were invisible across all three imaging layers. After AO correction, apical dendrites and their corresponding soma became clearly distinguishable with high resolution and SBR. Analysis of calcium transients revealed synchronized activity between apical dendrites and soma during whisker stimulation (Fig. S15e-g).

Next, we performed high-resolution imaging of neuronal dendrites and somas in the visual cortex during visual stimulation in awake mice. A liquid-crystal display (LCD) screen positioned 15 cm from the mouse’s eye displayed moving gratings at various angles, synchronized with image acquisition (see Methods). The improved imaging contrast and resolution after AO correction enabled precise localization of fine apical dendrites and somas, and extraction of calcium transients during visual stimulation (Fig. 3g, h, Supplementary Video S9). Analysis of neuronal responses to moving gratings identified direction-selective neurons and their dendrites (Fig. 3i-k), highlighting specialized neuronal functions^50,51^.

### Imaging of Microvascular Hemodynamics and Neurovascular Coupling (NVC)

The microvascular system, a crucial component of brain cortex, supplies oxygen and nutrients to neurons while removing waste, playing an essential role in brain function and health^52–54^. Microvascular hemodynamics closely relate to brain physiological state and are strongly influenced by anesthesia^55,56^. To study microvascular hemodynamics in natural conditions of brain, we imaged capillaries in awake mice using MD-FSS-2PM. After AO correction, fine capillary structures were well resolved (Fig. 4a, b), with line profiles demonstrating greatly improved resolution (Fig. 4c). The enhanced image quality enabled red blood cell (RBC) velocity measurements in brain capillaries of awake behaving mice using line-scan imaging, with velocity calculated from line scan kymographs using Radon transform algorithm^57^. We repeated these measurements after isoflurane anesthesia. Statistical analysis showed increased RBC velocity in the same capillaries during anesthesia (Fig. 4d, e), confirming isoflurane’s influence on cerebrovascular hemodynamics^55^.

**Fig. 4.**
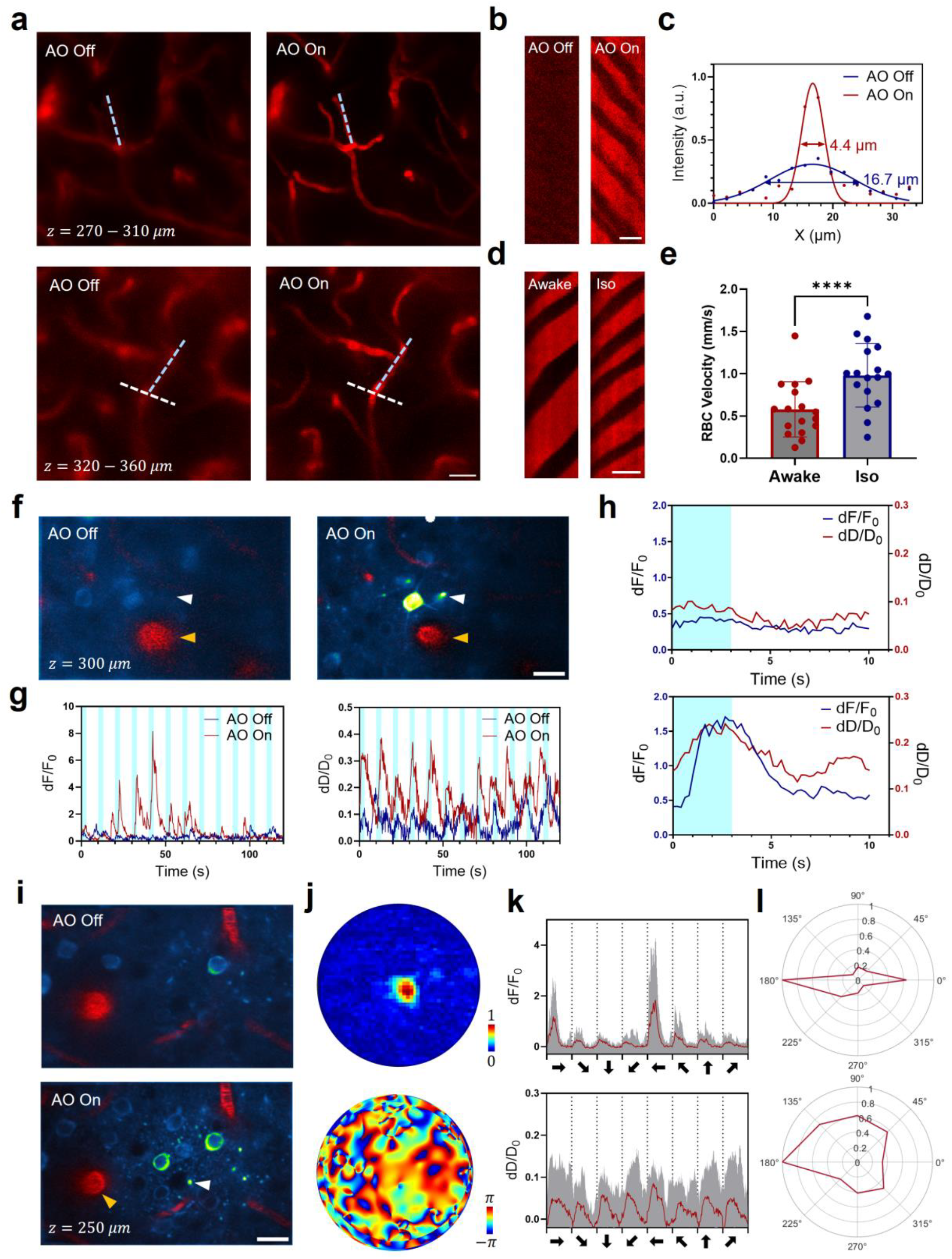
MD-FSS-2PM imaging of microvascular hemodynamics and neurovascular coupling in awake behaving mice. (a) MIP images of Texas Red-labelled blood vessels in the volume at depth of 270 − 310 *μm* (left) and 320 − 360 *μm* (right) below pia without (top) and with (bottom) AO correction on awake behaving mouse. (b) Kymographs of line scan images along the blue dashed lines in second row of (a), shown without (left) and with (right) AO correction. (c) Intensity profiles along the white dashed lines in the second row of (a) without and with AO correction. (d) Kymographs of line scan images of microvasculature on awake behaving mouse (left) and isoflurane-anesthetized mouse (right) along the blue dashed lines in the first row of (a) after AO correction. (e) Statistical analysis of red blood cell (RBC) velocity on awake behaving mouse and isoflurane-anesthetized mouse. Data were represented as mean ± SD. (Paired *t*-test, *α* = 0.05, N = 17 capillaries from 3 mice, ****p<0.0001); (f) Two-color merged images of STD projection images of GCaMP6s-labeled neurons over a 120 s recording and one frame images of Texas Red-labelled blood vessels during whisker stimulation at a depth of 300 *μm* below pia, shown without (left) and with (right) MD-FSS AO correction. (g) Left: Calcium transients extracted from the dendrites indicated by the white arrowhead in (f). Right: Diameter changes of penetrating arteriole (PA) marked by orange arrowhead in (f); blue stripe: whisker stimulation ON time. (i) Averaged traces of calcium transients of dendrites and PA diameter changes over 12 trials of whisker stimulation without (top) and with (bottom) AO correction; Blue stripe: whisker stimulation ON time. (i) Two-color merged images of STD projection images of GCaMP6s-labeled neurons over a 160 s recording and one frame images of Texas Red-labelled blood vessels during visual stimulation at depth of 250 *μm* below pia without (top) and with (bottom) MD-FSS AO correction. (j) Measured PSF amplitude (top) and AO correction phase pattern (bottom) in (i). (k) Top: Calcium transients in response to visual stimulation at different orientations of dendrites marked by white arrowhead in (i) after AO correction; Bottom: Changes in PA diameter in response to visual stimulation at different orientations marked by yellow arrowhead after AO correction; Gray curve: raw traces from multiple trials (N = 10) of random visual stimulation orientations; Red curve: averaged trace stimulated by different orientations; Gray dashed lines: timestamps of visual stimulation; Black arrowhead: orientation of visual stimulation. (l) Top: Orientation polar map of neuron dendrites indicated by white arrowhead in (i); Bottom: Orientation polar map of PA diameter marked by yellow arrowhead in (i). Awake: awake state; Iso: anesthetized by isoflurane. Scale bars: 20 *μm* in (a), (f) and (i); 10 *μm* in (b) and (d).

Neurovascular coupling (NVC), the modulation of cerebral blood flow (CBF) in response to regional neuronal activity, involves coordinated interactions between neurons and local vascular circulation crucial for supporting brain function^58–60^. To visualize neurovascular responses to stimulation in deep cortical regions, we performed multi-color imaging of neuronal activity and vascular dilation in the somatosensory cortex of awake mice. After AO correction, fine neuronal dendrites and penetrating arteriole (PA) outlines became clearly distinguishable (Fig. 4f, Supplementary Video S10), enabling accurate and sensitive monitoring of neuronal activity and local vascular dynamics. Both dendritic calcium transients and PA dilation traces showed significant enhancement with AO correction (Fig. 4g, h). The averaged traces during whisker stimulation revealed close coupling between dendritic neuronal activity and PA diameter (Fig. 4h). Due to poor imaging resolution, pre-AO correction traces showed significantly reduced sensitivity to stimulation in both neuronal activity and PA diameter changes compared to post-AO correction (Fig. 4h), demonstrating the necessity of MD-FSS AO correction for accurate NVC measurements. Furthermore, we conducted simultaneous multi-color imaging of neurons and blood vessels in the visual cortex during visual stimulation. AO correction greatly enhanced imaging resolution and intensity for both neurons and blood vessels (Fig. 4i, j, Supplementary Video S11). Analysis of synchronized dendritic calcium activity and PA diameter responses to directional visual stimulation revealed strong correlation between PA diameter and dendritic calcium responses (Fig. 4k). While the neuronal response was strongly direction-selective, PA dilation showed no significant directional selectivity in response to visual stimulation (Fig. 4l), suggesting that individual PAs may support multiple neurons with different directional preferences.

## Conclusions

In this study, we demonstrated that near noninvasive and high-resolution imaging in the awake, behaving mouse brain is critical for neuroscience research, as it avoids both brain inflammation and the confounding effects of anesthesia on cellular behavior and biological mechanisms. The major challenges that must be resolved are the aberrations and scattering from highly scattering media like the skull that severely distort excitation light, necessitating wavefront correction to restore image quality. Adaptive optics correction in awake mice presents an additional challenge, requiring rapid measurement to overcome motion artifacts from guidestars moving with brain tissue. We developed MD-FSS, a fast adaptive optics method for high-speed measurement of the aberrated complex-valued PSF, that effectively measures and corrects aberrations. This method enables high-resolution deep-tissue imaging through both thinned-skull and optical clearing windows in the awake mouse brain.

MD-FSS enables rapid and precise PSF measurements for correcting aberrations and scattering in awake mouse brains, achieving a 0.1s measurement time while maintaining the same SNR and signal enhancement as single beam methods under stable conditions. Unlike single beam approaches that fail due to motion artifacts (Fig. 1f, S8), MD-FSS ensures accurate full-sampled PSF reconstruction through stable amplitude and phase calibration of multiplexed weak scanning beams. The enhanced correction enables visualization of previously unresolvable features: microglial processes, red blood cells in finest microvessels, and calcium transients in neuronal dendrites, allowing simultaneous observation of microglia dynamics, microvascular circulation, neuronal activities in soma and dendrites, and neurovascular coupling in awake mice - critical measurements previously impossible without rapid aberration correction. While our current eight-beam implementation achieves 0.1s PSF measurements, the method can scale to more beams for faster acquisition, requiring only proportional increases in RF frequency channels and sampling rates.

## Methods

### Configuration of MD-FSS-2PM system

A detailed schematic diagram of the MD-FSS-2PM system is presented in Fig.S1. The excitation light source is a femtosecond laser with a bandwidth-limited pulse duration of 150 fs and a repetition rate of 80 MHz (Coherent, Axon 920). The internal dispersion compensation unit of the laser is adjusted to counteract system dispersion, compressing the pulse duration to 150 fs after the objective. The excitation laser is then directed through a pair of 4f relay lenses to expand the beam size, matching the aperture of an AOD (AA Optics, DTXS-400-920). The expanded beam is next directed to pass through the AOD. A high-power multi-frequency RF driver (MPDS8c, AA Opto-electronic) transmits an RF signal with multiple frequency components to the AOD, generating multiple 1^st^ order diffraction beams with different frequency offsets and angular separations. The 0^th^ beam and the 1^st^ order diffraction beams were utilized as the strong stationary beam and the weak scanning beams, respectively. The 0^th^ beam is then passed through a pair of 4f relay lenses (L1: *f* = 300 mm, L2: *f* = 300 mm) and an optical delay line DL is inserted into the arm of the 0^th^ beam after the 4f relay lenes to adjust the time delay between 0^th^ beam and 1^st^ beams for efficient interference. The scanning beams are directed through dual prisms PRM1 and PRM2 at Brewster angle of incidence for compensation of angular dispersion. A pair of 4f relay lenses (L3: *f* = 300 mm, L4: *f* = 300 mm) is placed after the dual prisms for expanding the beam to match the aperture size of an *x-y* galvo scanning mirror GM1xy, keeping the aperture plane of the AOD conjugated to the scanning mirror plane. The stationary beam and the scanning beams are then combined using a polarized beam splitter (PBS) and expanded by a pair of 4f relay lenses (L5: *f* = 40 mm, L6: *f* = 150 mm) to slightly overfill the aperture of the deformable mirror (DM97-15, Alpao), which is used to correct system aberrations based on a sensorless AO method^11^. A rotatable half-wave plate and a PBS are placed before the DM to control the intensity ratio and ensure that the polarization of the stationary beam matches that of the scanning beams for efficient interference. The DM is conjugated to a 3 mm *x*-galvanometric scan mirror GM2x using a pair of 4f relay lenses (L7: *f* = 200 mm, L8: *f* = 40 mm), which resizes the beam to fit the scanner’s aperture. The *x* and *y* scanners (GM2x, GM2y) are mutually conjugated by a pair of 4f relay lenes (L9, L10: *f* = 50 mm), each consisting of two achromatic doublets (*f* = 100 mm) in a Plössl configuration. The *y*-scanning mirror is conjugated to an electrically tunable lens (ETL) via the scan lens L11 (*f* = 36 mm) and the tube lens L12 (*f* = 200 mm). Subsequently, the ETL is conjugated to the back aperture of a high numerical aperture (NA) objective (XLPLN25XWMP2, Olympus) using a pair of achromatic doublets (L13: *f* = 222 mm and L14: *f* = 150 mm). A phase-only spatial light modulator (SLM) is positioned approximately 80 mm after L13 and is conjugated to the skull plane through a 4f optical system consisting of L14 and the objective lens. The beam diameter entering the back aperture of the objective is approximately 10 mm, resulting in an excitation numerical aperture (NA) of approximately 0.7. The objective is mounted on a motorized actuator to enable translation along the optical axis. The emitted fluorescence is collected by the same objective and directed to the photo-detection unit via dichroic mirror DCM1 (FF705-Di01-25×36, Semrock). A dichroic mirror DCM2 (F556-SDi01-25×36, FF495-Di03-25×36, Semrock) is placed after DCM1 to separate fluorescence with different color for dual-color imaging. A suitable band-pass filter, along with a short-pass filter, is placed before the photomultiplier tube (PMT; H5783, H7422, Hamamatsu) module to selectively detect the SHG/Texas Red and GFP/YFP/GCaMP6s signals. The current signal from the PMT is converted to a voltage signal using a transimpedance current amplifier (DHCPA-100, Femto; SR570, Stanford Research). For 2P imaging, only the stationary beam is directed to the sample while the scanning beams are blocked. Galvanometric scanning mirrors GM2x and GM2y are employed for lateral scanning. ETL is used to adjust the imaging depth. The excited fluorescence signal is digitized at a sampling rate of 2 MHz using a high-speed data acquisition card (PXIe-6363, NI Instrument). The microscope control software is custom written in C#.

### Multi-beam generation and multi-frequency phase modulation

The core principle of MD-FSS is based on generating multiple scanning beams with each beam undergoing linear phase modulation at a distinct frequency to ensure that adjacent beams are spatially separated with the line distance of the full-sampled PSF. An AOD can serve as a beam deflector, diffracting the input beam to the 1^st^ order angle of diffraction. A RF signal with components of multiple frequencies can be applied to the AOD, simultaneously generating multiple first-order diffraction beams, each corresponding to a specific RF frequency. The multi-frequency RF signal *V*_*RF*_ for *N* beam generation can be represented as:

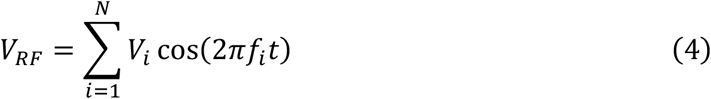

Where *f*_*i*_ is the *i*_*th*_ beam modulation frequency in the RF signal, *V*_*i*_ is the amplitude of frequency components *f*_*i*_. With the applied RF signal, the diffraction angle *θ*_*i*_ for each scanning beam can be expressed as:

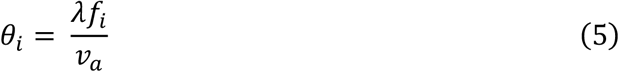

Here, *λ* represents the beam wavelength, and *υ*_*a*_ denotes the acoustic velocity in the acousto-optic crystal. Thus, the diffraction angle of each scanning beam is determined by its respective modulation frequency *f*_*i*_. Since the spatial distance between the foci of adjacent scanning beams must match the line distance of the full-sampled PSF, the frequency distance should be calculated based on the sampling distance in the focal field. The relationship between the diffraction angle and the sampling distance Δ*x* can be expressed as:

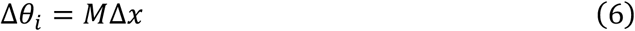

Where *M* represents the scaling factor between the AOD and the focal field, as determined by the magnification of the designed system. Linear phase modulation is automatically achieved by applying *V*_*RF*_, as the AOD functions as a frequency shifter, adding or reducing the optical frequency of the 1^st^ order diffracted beam by *ω*_*i*_ = 2*πf*_*i*_. Therefore, the phase modulation of each diffracted beam can be expressed as:

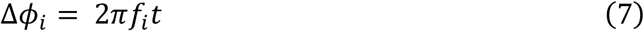

The wave function of each scanning beam can be expressed as:

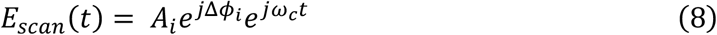

Where *A*_*i*_ is the amplitude of each scanning beam and *ω*_*c*_ is the optical frequency of the input beam. In this manner, each scanning beam is phase-modulated at a different frequency *f*_*i*_, and is equally shifted in space at the focal field.

### Multi-frequency phase and amplitude detection based on digital FFT

The detailed flowchart for digital FFT demodulation process is illustrated in Fig. S2. Since the acceptable RF frequency for AOD typically ranges in the tens of megahertz, it is impractical to demodulate the phase and amplitude at the exact modulation frequency. By leveraging the periodic properties of the two-photon femtosecond pulse train, the interference signal between the strong beam and the phase-modulated weak beams can be considered as being sampled at the repetition frequency of the femtosecond laser, which can be expressed as:

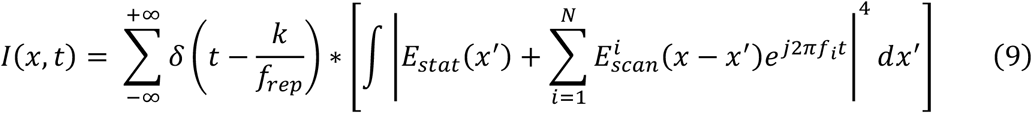

Where *f*_*rep*_ is the repetition frequency of the femtosecond pulse train, *f*_*i*_ is the modulation frequency of the *i*_*th*_ scanning beam and *N* denotes the number of multiplexed scanning beams. Using a high-order expansion approximation, the modulated fluorescence intensity of each scanning beam in frequency domain can be represented as the convolution of a periodic frequency comb with the corresponding modulation frequency peak after taking Fourier transform. Therefore, the available demodulation frequency for the *i*_*th*_ scanning beam can be expressed as:

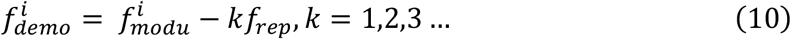

By setting the modulation frequency equal to the sum of the repetition frequency and a small offset frequency Δ*f*^*i*^, we have:

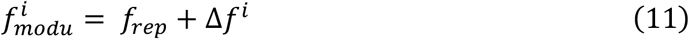

Thus, the demodulation frequency can be calculated as follows:

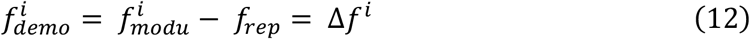

In this study, we selected the frequency offset list {Δ*f*^*i*^} starting with a value of 476 *kHz* and a frequency interval of 51 *kHz* to ensure equal spatial shift of 0.25 *μm* (Fig. S3). In this manner, the modulation frequency starts at 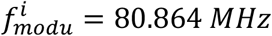 with an increase step of 51 *kHz*, given the laser repetition frequency *f*_*rep*_ = 80.388 *MHz*.

The modulated fluorescence signal is then sampled using a high-speed data acquisition card (PXIe-5170R, NI Instrument) and recorded on a computer for analysis. By applying Fast Fourier transform to the recorded fluorescence signal, the frequency spectrum at modulated frequencies *ω*_*i*_ can be represented as:

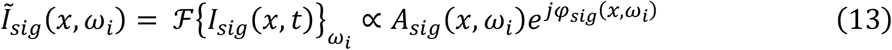

The mix of the RF modulation signals and the laser synchronization signal is recorded simultaneously as the reference for digital FFT demodulation. By applying Fast Fourier transform to the reference signal, the amplitude and phase at each demodulation frequency *ω*_*i*_ can be extracted as:

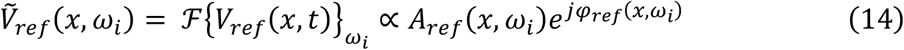

The complex electric field of the *i*_*th*_ scanning beam can then be computed as:

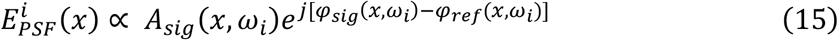

Since the recorded signal and reference contains information of all the scanning beams, the complex-valued PSF of each scanning beam can be calculated directly in the frequency domain.

### AOD Dispersion compensation

The AOD crystal introduces significant dispersion in both the stationary and scanning beams, which must be compensated for optimal performance in multi-beam interference. The temporal dispersion is counteracted by the build-in dispersion compensation unit of the laser. Since the diffraction angle of the 1^st^ order beam is linearly dependent on the wavelength and inversely proportional to the acoustic speed in the crystal, only the scanning beam experiences angular dispersion from the AOD. To compensate for the angular dispersion of the scanning beams, dual prisms are positioned after the AOD at the Brewster angle of incidence, with beam size magnification of 1 and maximum transmission (Fig. S4). The distance between the dual prisms and the AOD is minimized to prevent the introduction of additional temporal dispersion in the scanning beam. Each prism compensates for a portion of the total angular dispersion, ensuring that the final residual angular dispersion is minimized.

### Frequency calibration of RF driving signal on AOD

A calibration lookup table (LUT) is necessary to determine the exact frequency distance that corresponds to a specific spatial distance and the calibration method is illustrated in Fig. S3. Initially, a list of RF frequencies {*f*} near the most efficient working frequency *f*_*c*_ of the AOD is selected and applied to the AOD in sequence. The scanning beam is then utilized to image a singular fluorescent bead at each applied frequency using GM1xy, with the stationary beam blocked. For each applied frequency in the frequency list {*f*}, a fluorescence image is recorded. Finally, the center of mass of the fluorescent bead in each recorded image is calculated and fitted with a linear function *y* = *af* + *b*, given that *f* is the modulation frequency corresponding to each image. The spatial scan distance per RF driving frequency is determined by the coefficient *a*. In this work, we have *a* = −4.903 *μm*/*MHz*, resulting in a modulation frequency distance of 51 *kHz* between adjacent scanning beams with 0.25 *μm* spatial offset in the focal plane.

### Phase and amplitude calibration between multiplexed scanning beams

The relative phase and amplitude between the multiplexed scanning beams must be precisely calibrated for accurate reconstruction of a full-sampled PSF. First, the PSF of each scanning beam is acquired simultaneously without down-sampling when multi-frequency RF signal is applied. To determine the relative phase, the PSF of one scanning beam is selected as the reference PSF, and a list of phase offsets {Δ*φ*_*i*_} is sequentially added to phase map of other PSFs, represented as:

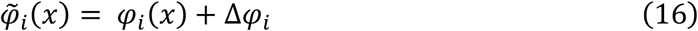

Where *φ*_*i*_ is the measured PSF phase map of the *i*_*th*_ scanning beam, and Δ*φ*_*i*_ is the phase offset added to the raw phase map. The correct phase offset Δ*φ*_*i*_ is determined by maximizing the correlation between the raw phase map *φ*_*i*_ and the adjusted phase map 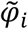. The calibration of amplitude is calculated by dividing the mean amplitude of reference PSF to that of each scanning beam PSF, expressed as:

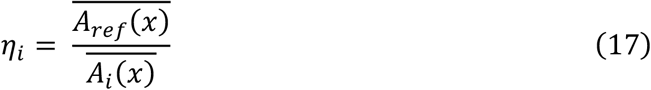

Using the amplitude ratio *η*_*i*_, the adjusted amplitude of each PSF is then calculated as:

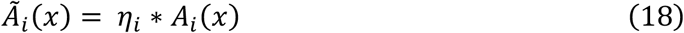

With the calibrated LUT of amplitude and phase, the complex-valued electric field of each scanning beam can be expressed as:

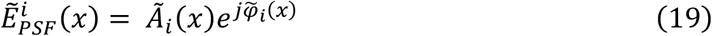

### Remote focusing and system aberration correction

To preserve the conjugation relationship between the SLM and the skull in deep tissue imaging, ETL is employed to adjust the position of the focal plane. A DM is utilized to correct the system aberration caused by misalignment and defocus of the ETL. In brief, the beam is initially focused on a Rhodamine 6G solution, and a series of remote focusing depths within the ETL range are adjusted sequentially. At each remote focusing depth, the fluorescent signal is used to measure system aberration according to a sensorless AO algorithm^11^. Once the aberration coefficient at each depth is determined, a LUT is created and used to generate the correction wavefront for different imaging depths. The wavefront between the LUT sampling points is calculated using linear interpolation of the wavefronts in the two nearest points^61^.

### Calibration of SLM conjugate position

The determination of the conjugation position of the SLM follows the previous method^17^. Briefly, a thin fluorescence layer is initially imaged using two-photon imaging, and the position of the objective is recorded as *z*_1_. Next, a camera is positioned in front of the SLM, and its position is adjusted to obtain a clear image of the SLM surface. The objective position is then finely adjusted using a motorized actuator until the image of the thin fluorescent layer is clearly visible on the camera, at which point the objective’s position is recorded as *z*_2_. The conjugate position of the SLM relative to the focal plane is then calculated as follows:

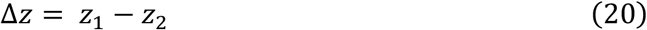

In our system, the SLM is conjugated to the plane with Δ*z* = 430 *μm*. For each *in-vivo* experiment, the focal plane is initially adjusted to the center of the mouse skull layer using the skull’s SHG signal as a guide, then it is moved downward by a distance of Δ*z*. Once the focus is adjusted to the target plane, the objective’s position is fixed, and remote focusing is employed to change the imaging depth.

### *In-vivo* imaging

For morphology imaging on awake behaving mouse, the mouse was initially placed on a treadmill, allowing it to run freely while its head was fixed to a rotational stage. The angle of the rotational stage was coarsely adjusted to flatten the skull based on a bright field image on a commercial camera and then finely tuned using the SHG image. After positioning the skull perpendicular to the excitation path, the conjugate plane was adjusted to align with the middle of the skull layer by modifying the axial position of the objective. Next, MD-FSS was utilized to measure the aberrated PSF using guidestars of microglia, neurons, and other sources, and the correction wavefront was calculated for application on the SLM. Finally, 2PF images with significantly enhanced resolution and signal intensity were obtained with the correction pattern applied on SLM.

### Animal preparation

All the animal experiments were conducted on adult mice (>=2 months old). Three transgenic mouse lines—*Thy1-YFP (Tg(Thy1-YFP)HJrs/J)*^62^, *Cx3Cr1-GFP (B6*.*129P2(Cg)-Cx3cr1tm1Litt/J)*^63^, *CaMKIIa-GCaMP6s. CaMKIIa-GCaMP6s* mice were generated by crossing *CaMKIIa-Cre (B6*.*Cg-Tg(Camk2a-cre)T29-1Stl/J)* mice with *tetO-GCaMP6s (B6;DBA-Tg(tetO-GCaMP6s)2Niell/J)* mice. All animal procedures were conducted in accordance with the Guidelines of the Animal Care Facility of the Hong Kong University of Science and Technology (HKUST) and were approved by the Animal Ethics Committee at HKUST.

### Drug administration method

Global systematic inflammation was induced by administration of LPS. Briefly, a single intraperitoneal injection of LPS (5 mg/kg) was repeated daily for three days prior to awake two-photon imaging. 100 *μL* Texas Red (2 mg/100 *μL*, Dextran, 70,000 MW, Thermo Fisher) was used for fluorescence labelling of blood vessels via retro-orbital injection. To induce anaesthesia during imaging, isoflurane was administrated with dose of 1%. Before thinned skull window and optical clearing window preparation, mice were anesthetized by intraperitoneal injection of xylazine (8.75 mg/kg) and ketamine (87.5 mg/kg).

### Thinned skull window preparation

The skull thinning procedure was slightly modified from a previous protocol^64^. After anaesthesia with xylazine and ketamine, the dorsal surface of the head was shaved, and the exposed skin was then cleaned and disinfected with 75% ethanol. An incision on the mid-line was made starting between the eyes to just caudal of the ears, and a lateral incision was made to the boundary of temporal muscle and skull. A drop of phosphate buffered saline (PBS) is applied on the exposed skull so that the periosteum above the skull can be lifted easily by forceps and removed by a curved surgical scissor. After the skull surface is dried up, tissue glue (3M Vetbond, Tissue Adhesive) is applied to the cut boundary of skin. A thin layer of C&B Metabond is then applied over the exposed skull. A customized headplate was then held firmly on the skull by applying a mixture of dental cement and cyanoacrylate glue. Then mouse was mounted on an angle adjuster (MAG-2, NARISHIGE, Japan) and a 0.5-mm carbon steel burr attached to a high-speed drill was used to thin a small circular region (diameter, 2-3 mm). Drill center is located using a stereotaxic apparatus (MingWangKeJi Ltd, Zhongshan, China) at 3.5 mm lateral and 1 mm posterior to the bregma for the somatosensory cortex, 2.5 mm lateral and 4 mm posterior to the bregma for the visual cortex. After majority of the middle spongy bone was removed, a microsurgical blade (no. 6961, Surgistar) was used to manually thin the skull to approximately 30-40 μm in thickness, which can be accurately measured using the SHG signal of the skull bone.

### Optical clearing intact skull window preparation

The procedure for optical clearing of the intact skull followed the established protocol^43^. The optical clearing solutions consist of S1 and S2. S1 is a saturated supernatant solution of 75% ethanol and urea at room temperature, while S2 is a high-concentration sodium dodecylbenzenesulfonate (SDBS) solution. To prepare S1, 75% ethanol (Sinopharm, China) was mixed with urea (Sinopharm, China). The mixture should be stirred continuously, then allowed to stand so that the supernatant can be collected for use. The volume-to-mass ratio of ethanol to urea is approximately 10:3. For S2, a 0.7 M NaOH solution is mixed with dodecylbenzenesulfonic acid (DDBSA, Aladdin, China) at a volume-to-mass ratio of 24:5, resulting in a pH of 7.2–8.0. The clearing maintenance reagent comprises of primer gel (Superior mixing clear gel, RyujiandMay, China), joint gel (K-303, Kafuter, China), and sealing gel (plated crystal antifouling sealing gel, Rainey, China). The LED light source (365nm) was used to make the gel fully solidify.

### Whisker stimulation method

The whisker stimulation method was slightly modified from the previous protocol^58^. In brief, a jet pipe was positioned near the mouse’s whisker and controlled by an electric gas valve. The air pressure was precisely adjusted to provide whisker stimulation with appropriate strength. The waveform of whisker stimulation adhered to the previous protocol^58^, featuring a pulse train with a duration of 3 seconds, a temporal frequency of 4 Hz, a pulse width of 100 ms, and an air pressure of 0.1 MPa. A custom-written MATLAB software was used to synchronize the image acquisition and whisker stimulation.

### Visual stimulation method

The visual stimulation method was slightly modified from the previous protocol^65^. In brief, an LCD screen was positioned at 30-45 degree relative to the mouse’s center line, 15 cm in front of the left eye. The LCD screen was covered with blackout material and filtered by a short-pass filter to prevent light contamination. A custom-written MATLAB program was used to generate moving gratings for directional visual stimulation, which was synchronized with imaging. The moving grating was generated with a spatial frequency of 0.05 *cc*/° and a temporal frequency of 2 Hz.

### Image analysis

The images were processed using MATLAB or ImageJ. To eliminate inter-frame motion artifacts, the images were registered using the TurboReg or StackReg plugin in ImageJ, as well as the NoRMCorre algorithm^66^ in MATLAB. Some images were processed using the Smooth function in ImageJ, which replaces each pixel’s value with that of its 3 × 3 neighboring pixels, or with the Gaussian Blur function, applying a sigma of 0.15–0.2 μm. The image pairs with AO Off and AO On correction were acquired and processed using the same parameters and displayed with consistent contrast. The microglia morphology and dynamics were analysed following previous work^21^. Regarding the calculation of the ramification index, process area changes, and Sholl analysis, the raw image was first normalized to a consistent saturation level and then smoothed using a Gaussian filter. The processed image was then binarized using the same threshold level. The ramification index (RI) was defined as the ratio of the perimeter to the area, normalized by the corresponding ratio for a circle with the same area, and was calculated from the binarized images. The area of bright pixels in the resulting mask was calculated for each image as the process area. Sholl analysis was conducted on the binary mask using the ImageJ plugin, Sholl Analysis 3.6.12. The calculation of calcium transients in neurons and the analysis of orientation selectivity of somas and dendrites were based on previous work^50,67,68^.

### Statistical analysis

Statistical analysis and data visualization were performed using GraphPad Prism 10 software. The data are presented as mean±SD, and *α* = 0.05 for all analyses. Data were analyzed using paired two-tailed t-test.

## Acknowledgement

The Hong Kong Research Grants Council through grants (16102122, 16102123, 16102421, 16102518, 16102920, T13-607/12R, T13-605/18W, C600217GF, C6001-19E, C6034-21G, T13-602/21N), the Innovation and Technology Commission (ITCPD/17-9), the Area of Excellence Scheme of the University Grants Committee (AoE/M-604/16, AOE/M-09/12) and the Hong Kong University of Science & Technology (HKUST) through grant 30 for 30 Research Initiative Scheme.

